# Read-Consistent Minimum Unique Substrings: A Parameter-Free, Linear-Time Framework for Genomic Sequence Representation

**DOI:** 10.64898/2026.02.28.708734

**Authors:** Andrews Frimpong Adu, Elliot Sarpong Menkah, Peter Amoako-Yirenkyi, Samson Pandam Salifu

## Abstract

Fixed-length k-mers have been the standard unit of genomic sequence representation for over two decades. However, they impose a uniform resolution on genomes whose complexity varies across loci. We introduce Minimum Unique Substrings (MUSs), variable-length sequence units defined by the local uniqueness structure of the genome rather than predefined parameters. We first extend MUS theory from single contiguous strings to fragmented sequencing reads by formalizing a definition of uniqueness that is consistent with these reads. Next, we present a linear-time extraction algorithm that runs in *O*(*n*) time using the generalized suffix tree. In this context, we introduce outpost nodes, topological anchors within the suffix tree that accurately localize MUS boundaries in fragmented sequencing reads. Finally, we empirically characterize the distributions of MUS lengths in *E. coli* K-12 and human chromosome 11. Our results demonstrate that MUS lengths naturally mirror genomic architectural complexity without the need for user-defined parameters. Notably, the MUS framework achieves 100% unique positional coverage with a mean length of only 36.08 bp. In contrast, fixed-length *k*=61 coverage reaches only 69.4%, despite being 1.69 times the MUS average. We show that increasing *k* from 21 to 61 triples the unique *k*-mer count from 2.35M to 6.86M. This *k*-paradox occurs because repetitive sequences are fragmented into spuriously unique tokens without improving true genomic resolution. MUSs escape this artifact entirely by adapting dynamically to local sequence complexity. These results establish MUSs as a biologically grounded, computationally tractable foundation for parameter-free genome assembly, repeat characterization, and alignment-free genomics.

## 1. Introduction

The rapid growth of sequencing data has significantly increased the demand for more efficient and scalable methods to represent genomic sequences [5, 6, 12, 32]. For over two decades, fixed-length *k*-mers–contiguous nucleotide sequences of length *k*–have served as the dominant representational unit in bioinformatics. They underpin critical applications, including genome assembly, variant detection, metagenomics, and comparative genomics [1, 16, 18, 19, 26, 27].

While their simplicity has facilitated considerable algorithmic advancements, fixed-length *k*-mers impose a uniform resolution on genomes that are inherently heterogeneous. A single *k* value must strike a balance between sensitivity and specificity across diverse genomic regions. As a result, small *k* values collapse distinct loci in repetitive regions into indistinguishable sequences, whereas large *k* values introduce fragmentation in unique regions [4, 22, 29, 33]. Because no single fixed *k* provides optimal resolution for an entire genome, a variety of adaptive methods have been proposed including iterative and multi-*k* approaches [3, 25], minimizer-based sparsification, context-dependent seeds, gapped k-mers, and substring sampling [7, 23, 28, 31]. These methods offer practical improvements but do not resolve the underlying fixed-length limitation. The representational unit remains externally defined by an arbitrary fixed-length parameter, rather than by the sequence’s intrinsic complexity.

Thus, the question of what constitutes a natural, data-driven unit of genomic resolution remains unanswered. Minimum Unique Substrings (MUSs)—substrings that occur exactly once in a sequence and cannot be shortened on either side without losing uniqueness—provide a principled solution to this problem. MUSs have been studied extensively within the combinatorics-on-words community [11, 17, 21, 24]. Their duality with maximum repeats establishes a direct, symmetric relationship between the unique and repetitive regions of a genomic sequence. Specifically, every MUS lies at the boundary of at least one repeat, and every repeat implies the existence of flanking MUSs [13]. This duality makes MUSs the natural unit for delineating repeat boundaries. However, prior theoretical work has been restricted to single contiguous strings, rendering it inapplicable to the de novo sequence analysis setting where the genome is unknown and the input consists of fragmented, overlapping sequencing reads.

We bridge this gap by introducing a read-consistent definition of MUS uniqueness—formalizing uniqueness with respect to a collection of reads rather than a complete genome. We then present a linear-time extraction algorithm that operationalizes this definition without a reference sequence. This algorithm introduces outpost nodes, a novel construction within the Generalized Suffix Tree that functions as topological anchors for MUS boundary localization across fragmented read data. The resulting framework achieves 100% positional uniqueness with substantially fewer and shorter tokens than any fixed-*k* representation. This provides both the empirical and theoretical foundations for graph-based genomic analysis and sequence indexing, in which representational units are data-intrinsic rather than parameter-defined. The contributions of this work are: (1) a formal, read-consistent extension of MUS theory to fragmented sequencing reads; (2) a linear-time 𝒪(*n*) extraction algorithm utilizing outpost nodes within a Generalized Suffix Tree; and (3) an empirical characterization of MUS distributions across diverse genomes, showing that MUS lengths naturally encode genomic architectural complexity.

## 2. Conceptual and Theoretical Framework

### 2.1 Preliminaries: Strings, Suffix Tries, and Suffix Trees

Let Σ = *{A, C, T, G}* denote an alphabet of size *σ*, and let *D* be a string over Σ, represented as *D* = *D*[1]*D*[2] … *D*[*n*]. We define *n* = |*D*| as the length of *D* and *D*[*i*] (1 *≤ i ≤ n*) as the *i*-th character in *D*. A substring of *D* is defined as *D*[*i* : *j*] (1 *≤ i ≤ j ≤ n*) spanning positions *i* and *j*, i.e., *D*[*i* : *j*] = *D*[*i*]*D*[*i* + 1] … *D*[*j*], which is empty if *i > j*. The reverse of *D* is denoted *D*_*r*_ = *D*[*n*] … *D*[1]. The *suffix* of *D* beginning at position *i* is suf[*i* : *n*] = *D*[*i*]*D*[*i* + 1] … *D*[*n*], and the *prefix* of *D* ending at position *i* is *D*[1 : *i*] = *D*[1]*D*[2] … *D*[*i*]. For a substring *u*, the number of its occurrences in *D* is given by *occ*_*D*_(*u*) = |*{ i* | *D*[*i* : *i* + |*u*| − 1] = *u }*|. The *suffix trie* of a string *D* is a rooted directed tree that contains all suffixes of *D*. Each node corresponds to a distinct substring of *D*, represented by the concatenation of edge labels from the root to that node. For substrings *D*_1_ and *D*_2_ of *D*, a directed edge (*D*_1_, *D*_2_) labeled by a character *α* exists if *D*_2_ = *D*_1_*α*. The *suffix tree S*(*D*) is a compact representation of the suffix trie of *D*. It merges non-branching paths into single edges, each labeled by a substring of *D*. By storing edge labels as pairs of indices referencing *D, S*(*D*) achieves linear space complexity. An *implicit suffix tree* is derived from *S*(*D*) by removing terminal symbols, unlabeled edges, and nodes with fewer than two children. A *suffix link* in *S*(*D*) connects internal nodes. Thus, if an internal node represents a substring *D*_1_ = *αD*_2_, its suffix link points to the node corresponding to *D*_2_. These links exist only between internal nodes, and if *α* is empty, the link points to the root.

The *generalized suffix tree* 𝒯(ℛ) for a collection of strings ℛ = *{D*_1_, *D*_2_, …, *D*_*k*_*}* is a compact trie that represents all suffixes of every string in ℛ. Each string *D*_*k*_ is typically terminated with a unique end symbol $_*k*_ to ensure that no suffix of one string is a prefix of another.

### 2.2 Minimum Unique Substrings (MUS)

Let *w* be a non-empty substring of *D* such that *occ*_*D*_(*w*) = *{ i* | 1 *≤ i ≤* |*D*| − |*w*| + 1, *D*[*i* : *i* + |*w*| − 1] = *w }*. We say that *w* is *unique* in *D* if it occurs exactly once, that is, |*occ*_*D*_(*w*)| = 1.

#### Definition 2.1

(Left and Right MUS). A unique substring *w* = *D*[*i* : *j*] is called a *left minimum unique substring* (`LMUS`) of *D* if it satisfies: |*occ*_*D*_(*w*)| = 1 and |*occ*_*D*_(*w*[1 : *k*])| *>* 1, ∀ 1 *≤ k <* |*w*|. Conversely, *w* = *D*[*i* : *j*] is a *right minimum unique substring* (`RMUS`) of *D* if: |*occ*_*D*_(*w*)| = 1 and |*occ*_*D*_(*w*[*k* : |*w*|])| *>* 1, ∀ 2 *≤ k ≤* |*w*|. An illustration of both types is provided in Figure 1.

**Figure 1:**
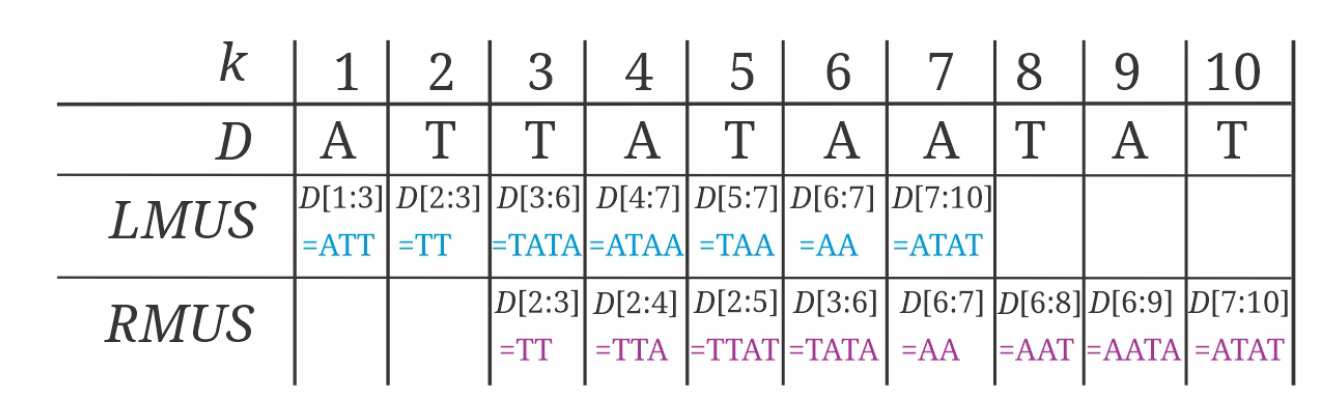
The `LMUS` and `RMUS` in *D* = *ATTATAATAT* for each position *k*. For position *k* = 4 we have *D*[4 : 7] = `ATAA`. Its proper suffix *D*[4 : 6] = `ATA` occurs more than once in *D* (i.e., it is a repeat), so ATAA cannot be shortened on the right while remaining unique. Hence the unique substrings that begin at index 4 include *D*[4 : 7] = `ATAA`, *D*[4 : 8] = `ATAAT` and *D*[4 : 10] = `ATAATAT`, and the shortest of these is *D*[4 : 7], which is therefore an `RMUS` starting at 4. Applying the LMUS/RMUS definitions across all positions yields the minimum unique substrings ℳ = {*D*[2 : 3] = `TT`, *D*[3 : 6] = `TATA`, *D*[6 : 7] = `AA`, *D*[7 : 10] = `ATAT` }. Each element of is ℳ the shortest unique substring at its start position (RMUS) or end position (LMUS), depending on context.

#### Definition 2.2

(Minimum Unique Substrings (MUS)). A unique substring *D*[*i* : *j*] is said to be a *minimum unique substring* (MUS) if it satisfies both the `LMUS` and `RMUS` conditions at indices *i* and *j*, respectively. In other words, *D*[*i* : *j*] is unique and cannot be shortened on either side while remaining unique. Formally, the set of intervals corresponding to the MUSs of *D*, denoted by ℳ_*D*_, is defined as:

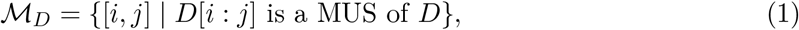

and the set of all MUSs in *D* is referred to as the `anchor set`, *A*_*w*_(*D*).

### 2.3 Uniqueness with respect to a set of reads

For a single, contiguous string *D*, the notion of uniqueness is unambiguous. Thus, a substring *w* is unique in *D* if and only if it occurs at exactly one position, that is, |occ_*D*_(*w*)| = 1. This definition supports a direct computational test and underpins the MUS framework of Section 2.2. In the context of genomic sequence analysis, the input is a collection of short/long, overlapping sequencing reads 𝒟 = *{d*_1_, *d*_2_, …, *d*_*m*_*}*, each of which is a fragment of a genome *D*. The notion of uniqueness is therefore extended through the concept of *consistency*, allowing algorithmic verification across multiple sequences. Although a substring *w* may appear in several strings, it may still correspond to a single unique location within a *Superstring*(*w*) (the minimal superstring that contains all strings in 𝒟 where *w* occurs). Consistency is therefore formalized as the following definition:

#### Definition 2.3.

A substring *w* is *consistent* if it occurs at most once in each *d*_*i*_ ∈ 𝒟, and each such *d*_*i*_ is a substring of *Superstring*(*w*) [15].

This definition then leads to the following proposition:

#### Proposition 1.

A word *w* is consistent if and only if:

1. It appears at most once within each read.
2. The reads containing *w* can be uniquely assembled into the shortest possible superstring, `Superstring(w)`, encompassing all of them.

Thus, Condition 1 ensures no ambiguous placement within any single read whiles Condition 2 ensures those reads agree on a single genome locus.

### 2.4 Maximum Repeats (MR)

A substring *w* is called a *repeat* if *occ*_*D*_(*w*) *≥* 2. A repeat is said to be *right-maximal* (`RMR`) if *D*[*i* : *j*] occurs at least twice in *D*, and its right extension *D*[*i* : *j* + 1], if defined, does not. Similarly, *w* is *left-maximal* (`LMR`) if *D*[*i* : *j*] occurs at least twice, and its left extension *D*[*i −* 1 : *j*], if defined, does not. A *maximum repeat* (MR) is therefore both `LMR` and `RMR.` Thus, MR cannot be extended in either direction without losing its repetitiveness. For instance, in Figure 1, *D*[4 : 6] = `ATA` and *D*[7 : 9] = `ATA` are MRs because they cannot be extended left or right while remaining repeats.

### 2.5 Duality Between MUSs and MRs

The duality principle establishes a direct, symmetric relationship between unique and repetitive regions of a sequence [13]. A MUS is a unique substring that extends from a MR, with each MUS arising at the boundary of a repeat.

#### Theorem 2

(Duality of MUSs and MRs). *For any genomic sequence D, a duality exists between MUSs and MRs such that:*

1. *Every MUS lies at the boundary of at least one repeat*.
2. *Every repeat implies the existence of at least one MUS in its vicinity*.

Intuitively, Theorem 2 shows that MUSs act as sequence anchors between repetitive regions. Notably: (1) If an MR begins at the start of the sequence (*i* = 1), then no left-extending MUS *D*[*i −* 1 : *l*] exists. (2) Similarly, if an MR ends at the last character (*j* = *n*), then no right-extending MUS *D*[*k* : *j* + 1] exists. Consequently (illustrated in Figure 2 as a result of Theorem 2):

**Figure 2:**
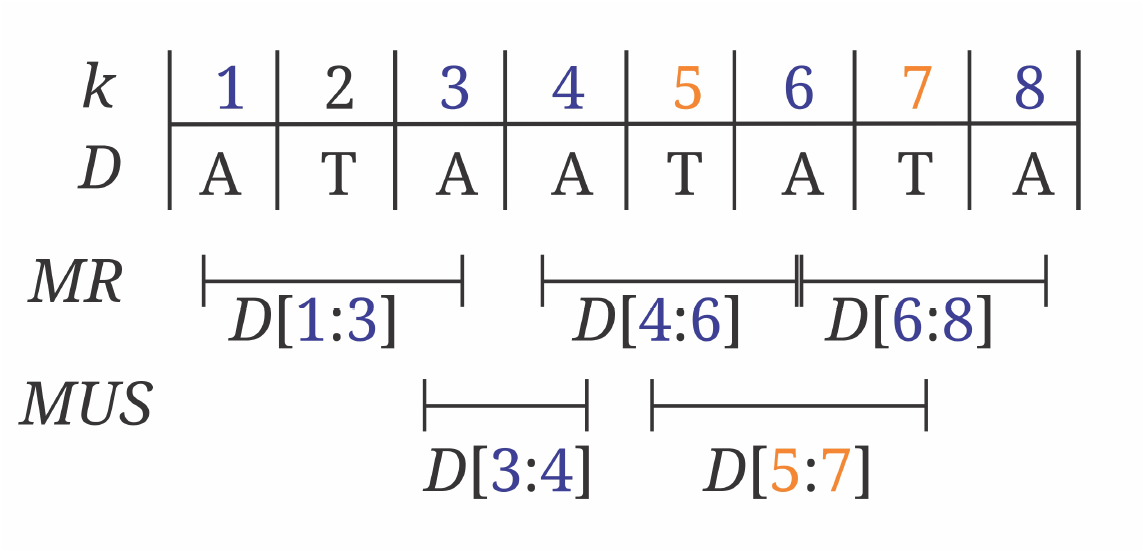
*(a)* The MUS *D*[*i* : *j*] implies the existence of MRs *D*[*k* : *j* − 1] and *D*[*i* + 1 : *l*] [13]. For example, for MUS *D*[5 : 7], we find the MRs *D*[4 : 6] and *D*[6 : 8], giving *k* = 4 and *l* = 8. *(b)* Conversely, the MR *D*[*i* : *j*] indicates the presence of MUSs *D*[*i* −1 : *l*] and *D*[*k* : *j* + 1]. For instance, with MR *D*[4 : 6], we identify the MUSs *D*[3 : 4] and *D*[5 : 7], yielding *k* = 5 and *l* = 4.

- For any MUS *D*[*i* : *j*], 1 *≤ i < j ≤ n*, there exist indices *k ≤ j −* 1 and *l ≥ i* + 1 such that *D*[*k* : *j −* 1] and *D*[*i* + 1 : *l*] are MRs.
- For any MR *D*[*i* : *j*], 1 *≤ i < j ≤ n*: (a) if *i ≥* 2, then there exists *l ≤ j* such that *D*[*i−*1 : *l*] is a MUS; (b) if *j ≤ n −* 1, then there exists *k ≥ i* such that *D*[*k* : *j* + 1] is a MUS.

### 2.6 Structural Properties

#### Monotonicity

If *D*[*i* : *j*] is a MUS of *D*, then any superstring *D*[*i*′ : *j*′] such that *i*′ *≤ i* and *j*′ *≥ j* and *D*[*i*′ : *j*′] *≥ D*[*i* : *j*] is also unique in *D*. Once uniqueness is achieved, it persists as you grow the substring outward.

#### Coverage of a position

For a position *t* in *D* (1 *≤ t ≤* |*D*|), the set of MUSs containing *t* is given as: *MUS*(*t*) = *{m* ∈ *MUS*(*D*)| ∃*i, j* s.t. *m* = *D*[*i* : *j*] and *i ≤ t ≤ j}*.

#### Existence

For the string *D* of length |*D*|, there exists at least one MUS in *D* (MUS(*D*)≠∅). Thus, *D* itself is unique, therefore, either at least one substring of *D* or *D* itself is a MUS.

##### Theorem 3

(Coverage bounds). *For any position t in D, with* 1 *≤ t ≤* |*D*|, *the number of MUSs covering t satisfies* 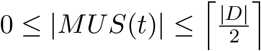, *where MUS*(*t*) *is the set of all MUSs that contain t. Both the lower bound* 0 *and the upper bound* ⌈|*D*|/2⌉ *are tight*.

Theorem 3 quantifies the number of MUSs that span a given position, demonstrating that MUSs may overlap. These properties imply that MUSs constitute an ordered, sparse set of anchors that delineate transitions between repetitive and unique regions of the genome.

## 3. Algorithmic Framework for MUS Extraction

The duality in Section 2.5 helps in identifying MUSs as boundaries between neighboring MRs. Given a genome of length *n*, MUSs are computed using a linear-time generalized suffix tree *T* (*R*) built with the Ukkonen’s algorithm [9]. This method employs the concept of outposts, which are specific suffix tree nodes marking where repeats in the genome transition to unique sequences. Every outpost is classified as either left or right boundary of at least one MUS. Figure 3 outlines the process of MUS identification: (1) Construct *T* (*R*) online, (2) Find right and left outposts, (3) Derive the corresponding MUS intervals, and (4) Store them in an ordered array for downstream analysis.

**Figure 3:**
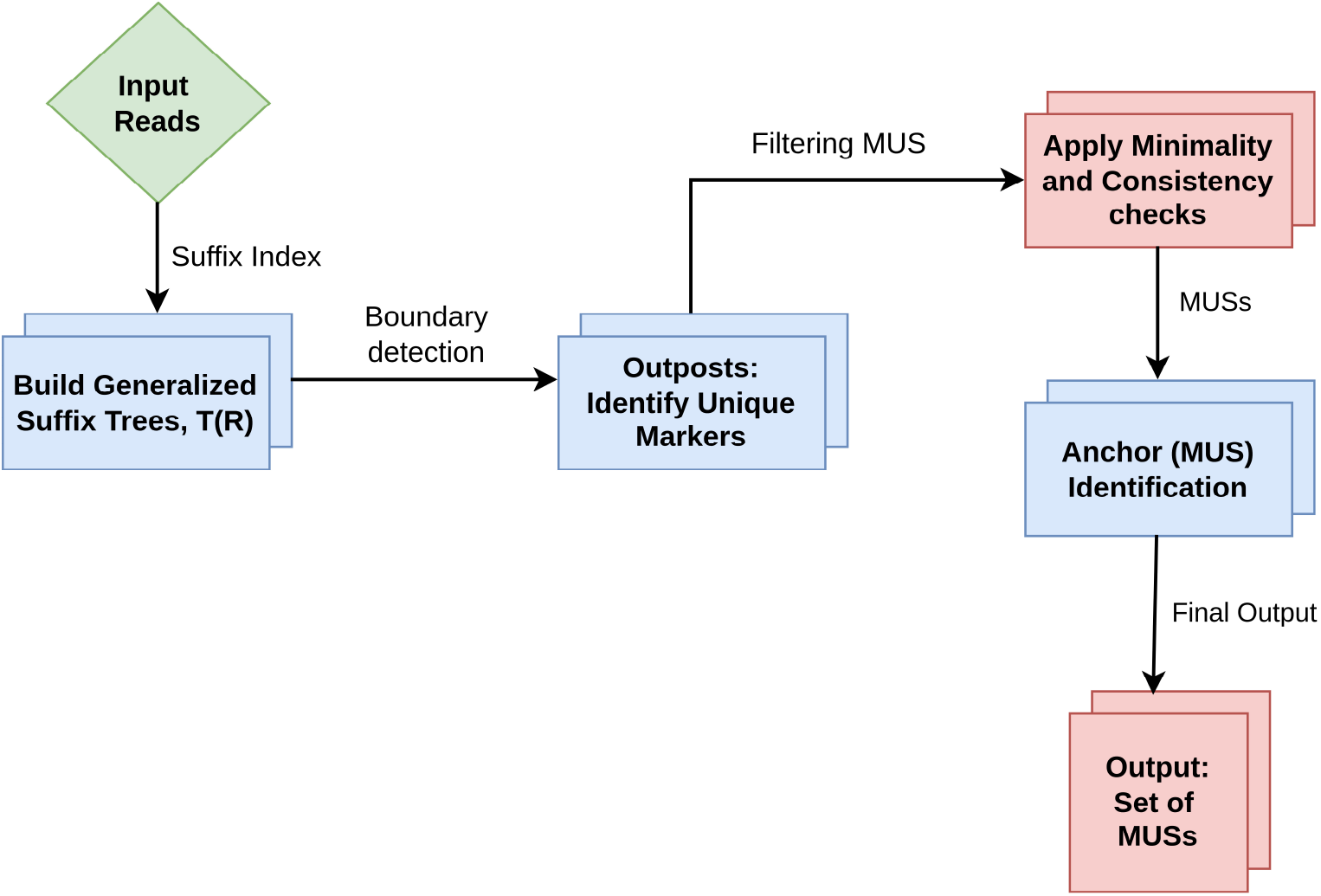
**MUSs extraction outline** from generalized suffix tree 𝒯 (ℛ) construction from a set of input reads, ℛ MUSs extraction.

### 3.1 Suffix Tree construction

Let ℛ = *{read*_1_, *read*_2_, …, *read*_*m*_*}* be a substring-free set of *m* reads, where each *read*_*k*_ = *r*_1_*r*_2_ … *r*_*n*_ has length |*read*_*k*_|. For each *read*_*k*_ (1 *≤ k ≤ m*), let *S*_*k*_ denote its suffix tree, and *S*_0_ the empty suffix tree (consisting only of the root). A unique terminal symbol $_*k*_ is appended to each read to form *read*_*k*_$_*k*_. The suffix set of a read *S* is defined as Suf[*S*] = *{S*[*i* : |*S*|] | 0 *≤ i ≤* |*S*|*}. Ukkonen’s algorithm* builds a suffix tree for a given *read* of length *N* in 𝒪(*N* ) time. The algorithm works in *N* phases, one for each character in the string. In each phase, the algorithm updates the suffix tree through a series of extensions. It achieves its linear time performance by sing *suffix links, implicit suffix trees*, and different update rules. The process begins by building a suffix tree *S* for the first read, *read*_1_. For every subsequent read, *read*_*k*_ (where *k ≥* 2), the algorithm begins by traversing the current suffix tree from the root. It tries to match a prefix of *read*_*k*_. If the first *i* characters of *read*_*k*_ match a path in the tree, this means all suffixes of *read*_*k*_ prefix (up to character *i−* 1) are already part of the tree (Proposition 4). Thus, the first *i* phases of Ukkonen’s algorithm for *read*_*k*_ are already done within the existing tree. The algorithm then picks up from phase *i* to add the remaining suffixes of *read*_*k*_. This method is shown in Algorithm 1. By repeating this for each part of the string, we end up with a tree that efficiently shows all suffixes of every input read, which is the generalized suffix tree 𝒯 (ℛ).

#### Proposition 4

(*Implicit Prefix Completion*). If the longest prefix of *read*_*k*_, say *read*_*k*_[0 : *i −* 1], matches a path in the already constructed tree *S*_*k−*1_, then all suffixes of this prefix, *read*_*k*_[*j* : *i−*1] for 0 *≤ j < i*, are inherently represented within *S*_*k−*1_. This effectively means that the first *i* phases of the Ukkonen’s algorithm for *read*_*k*_ have been satisfied by the existing tree structure.

#### Algorithm 1

Incremental Generalized Suffix Tree Construction

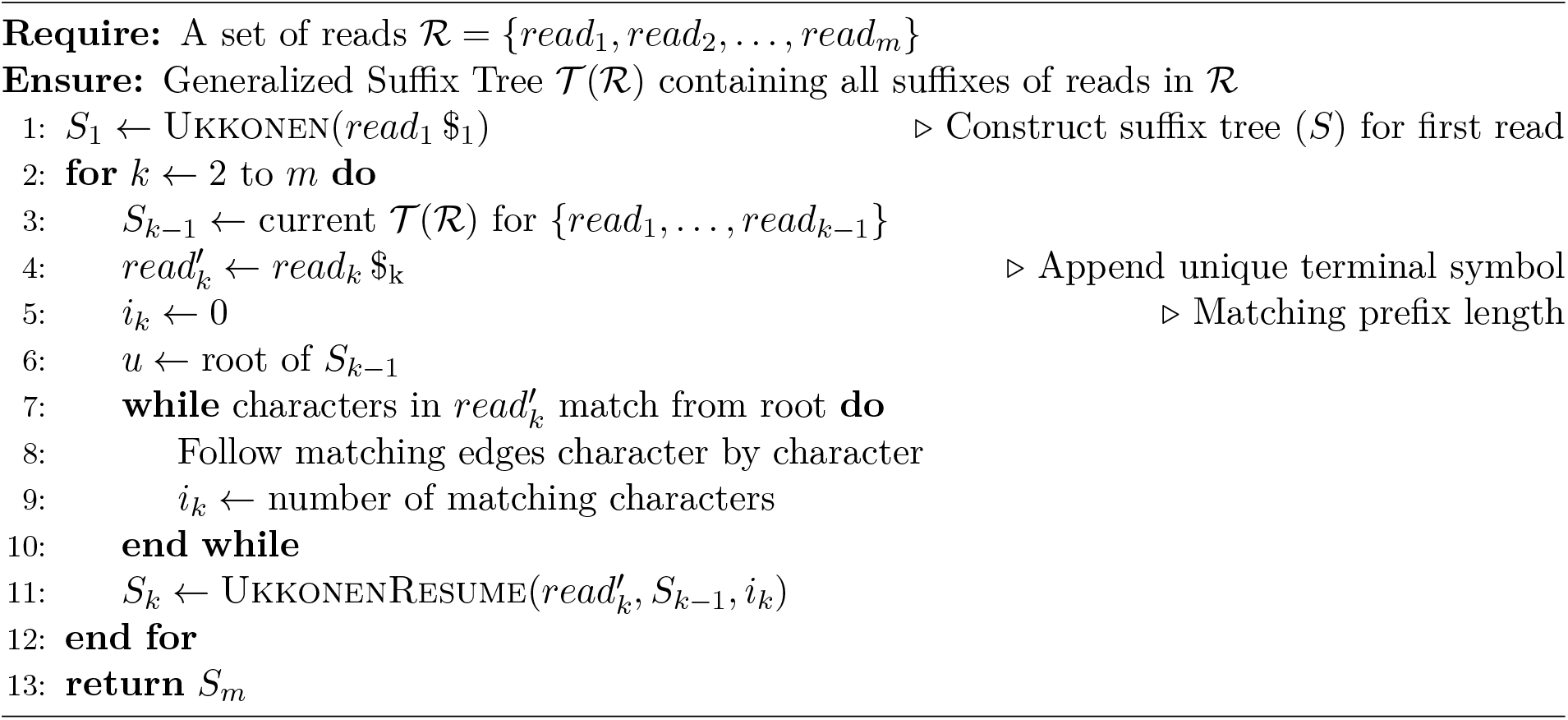

### 3.2 Outposts Identification

Let’s define Suf[*k, i*] as the suffix of *read*_*k*_$_*k*_ that starts at position *i* in 𝒯 (ℛ). This means Suf[*k, i*] = *read*_*k*_[*i* : |*read*_*k*_|]$_*k*_. Every Suf[*k, i*] has a unique path from the root to a leaf in 𝒯 (ℛ). We write *w*(*k, i, j*) as a substring of *read*_*k*_ from index *i* to *j*. The collection of all such *w* from ℛ is 𝒲(ℛ). All the suffixes together are written as 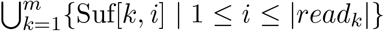. The label of a node *v*, written as `label`(*v*), is formed by joining the labels on the edges from the root to *v*. For an edge (*u, v*) in 𝒯 (ℛ), the substring labeling it is `label`(*u, v*).

#### Definition 3.1

(Trivial and Non-Trivial Edges). An *edge* (*u, v*) in 𝒯 (ℛ) is *trivial* if its label is a single unique termination character $_*k*_, i.e., label(*u, v*) = $_*k*_ for 1 *≤ k ≤ m*. Otherwise, if the label corresponds to a non-terminal substring *read*_*k*_[*i* : *j*], it is a *non-trivial edge*.

#### Definition 3.2

(Junction Node). A node *v* in 𝒯 (ℛ)is a *junction node* if it has at least two outgoing non-trivial edges, i.e., deg_Σ_(*v*) *≥* 2, where deg_Σ_(*v*) counts the number of such edges.

A junction node thus marks a branching point where multiple suffixes share a common prefix, with outgoing edges leading to distinct suffix extensions. Given a subset 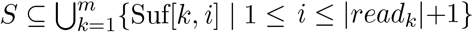, the suffixes in *S* originate from distinct reads if |*S*| = |*{k* | ∃ *i* such that Suf[*k, i*] ∈ *S}*|. For an edge (*u, v*), all suffixes (leaves) in the subtree rooted at *v* are said to occur *after* node *v* in 𝒯 (ℛ). These definitions form the basis for identifying MUSs during suffix tree traversal, using the concept of an *outpost*.

#### 3.2.1 Outposts

##### Definition 3.3

(Outpost). An outpost refers to a unique prefix of a suffix. This prefix is characterized by the fact that the rest of the suffix forms the complete label of the final edge that connects to the leaf node of that suffix (see Figure 4). To be precise, a word *w*(*i, j*) qualifies as an outpost if it serves as a prefix for some suffix Suf[*i*], and the unshared segment of that suffix corresponds to the label of a meaningful edge within the suffix tree.

**Figure 4:**
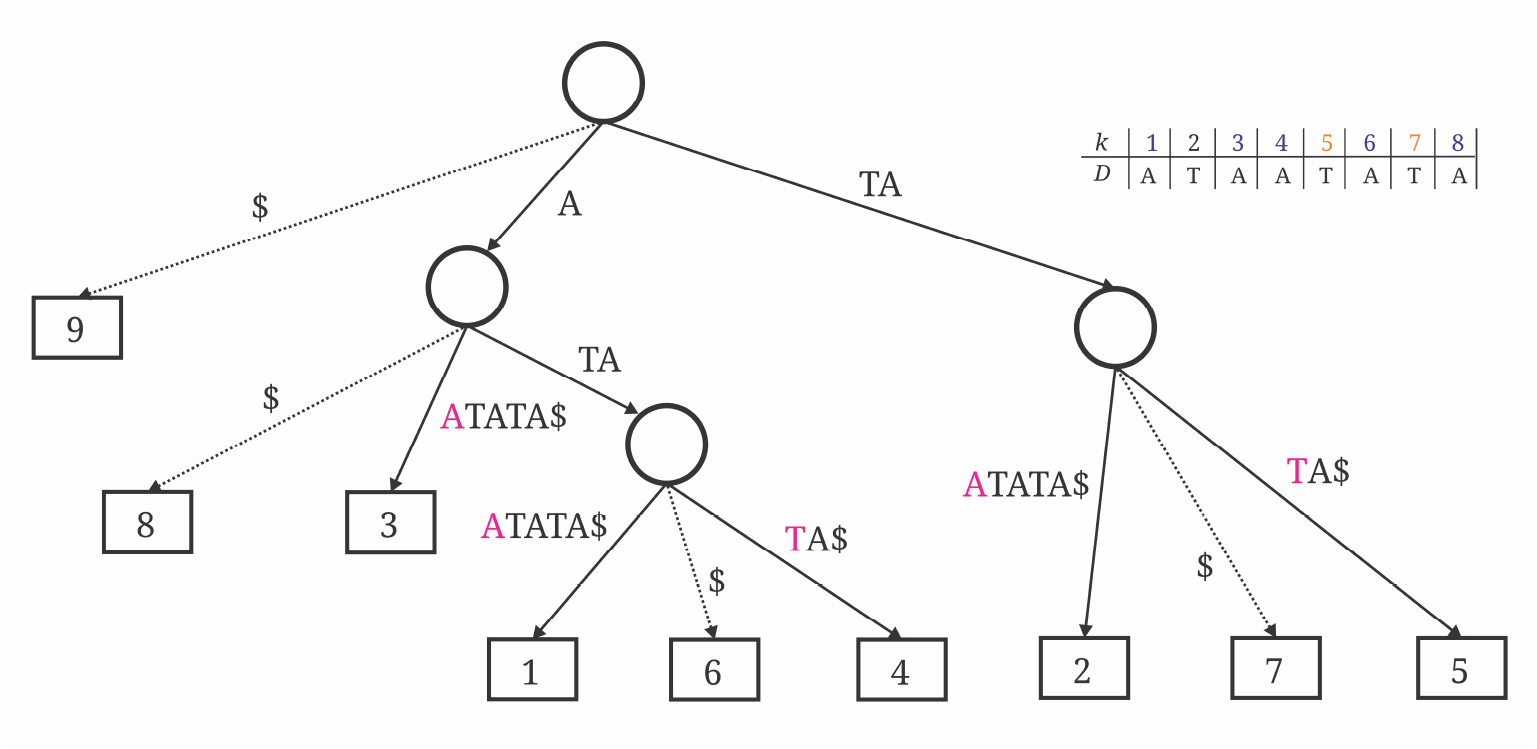
The suffix tree for a read *D* =ATAATATA. Trivial edges are shown in dotted lines and ends in only $. Branching nodes include A and TA ending in 2 and 5, 1 and 4, and 3. Outposts are ATAA, AA, ATAT, TAA, TAT.

Outposts are categorized as `right or left` and are identified along the path from the root to a leaf in 𝒯 (ℛ) corresponding to Suf[*k, i*] = *w*(*k, i*, |*read*_*k*_|)$_*k*_. Along this path, we locate the first edge (*u, v*) satisfying the following conditions:

1. All suffixes in the subtree rooted at *v* originate from `distinct reads`, and
2. The subtree under *v* is `not a junction.`

#### 3.2.2 Right and Left Outposts

If such an edge (*u, v*) exists, the substring up to node *v* defines a *right outpost*, labeled `label`(*u, v*) = *w*(*k, j, l*) if *l <* |*read*_*k*_|, or *w*(*k, j*, |*read*_*k*_|)$_*k*_ otherwise. Formally,

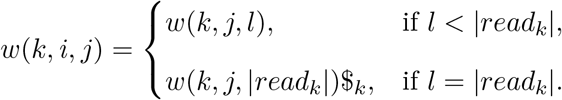

We denote Right_*k*_(*i*) = *j* as the boundary of the outpost, or Right_*k*_(*i*) = |*read*_*k*_| + 1 if no outpost exists for the suffix starting at position *i*. The procedure for detecting *Right*_*k*_(*i*) = *j* (identifying the end *j* of a word *w*(*k, i, j*) that starts at *i*) is depicted in Algorithm 2.

##### Algorithm 2

Find Right Outpost in *T* (*R*), Right_k

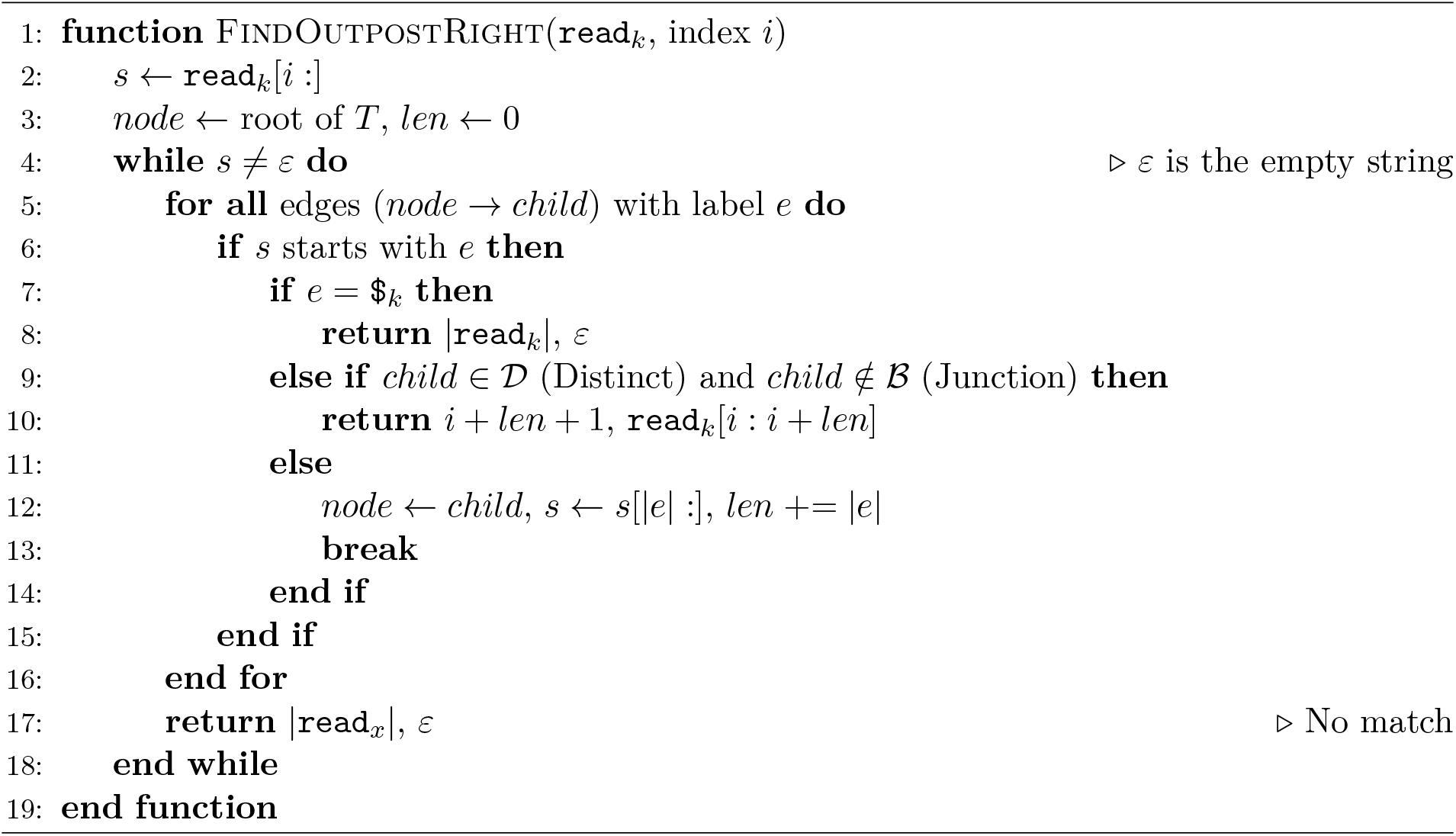

Symmetrically, *left outposts* can be identified by reversing the reads in *R*, denoted as 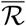, with each reversed read represented as 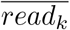 for 1 *≤ k ≤ m*. A left outpost is defined during the root-to-leaf traversal of the suffix tree constructed from 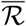, where the first edge (*u, v*) along the path satisfies the conditions (1) and (2) in 3.2.1. For such an edge, the label `label`(*u, v*) is given by

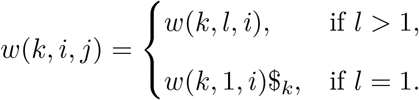

We define the boundary Left_*k*_(*j*) = *i*; if no such edge exists, then Left_*k*_(*j*) = 0. The procedure for detecting Left_*k*_(*j*) = *i*, which identifies the starting index *i* of a substring *w*(*k, i, j*) ending at *j*, is illustrated in Algorithm 3.

##### Algorithm 3

Find Left Outpost in *𝒯* (*ℛ*), `Left_k`

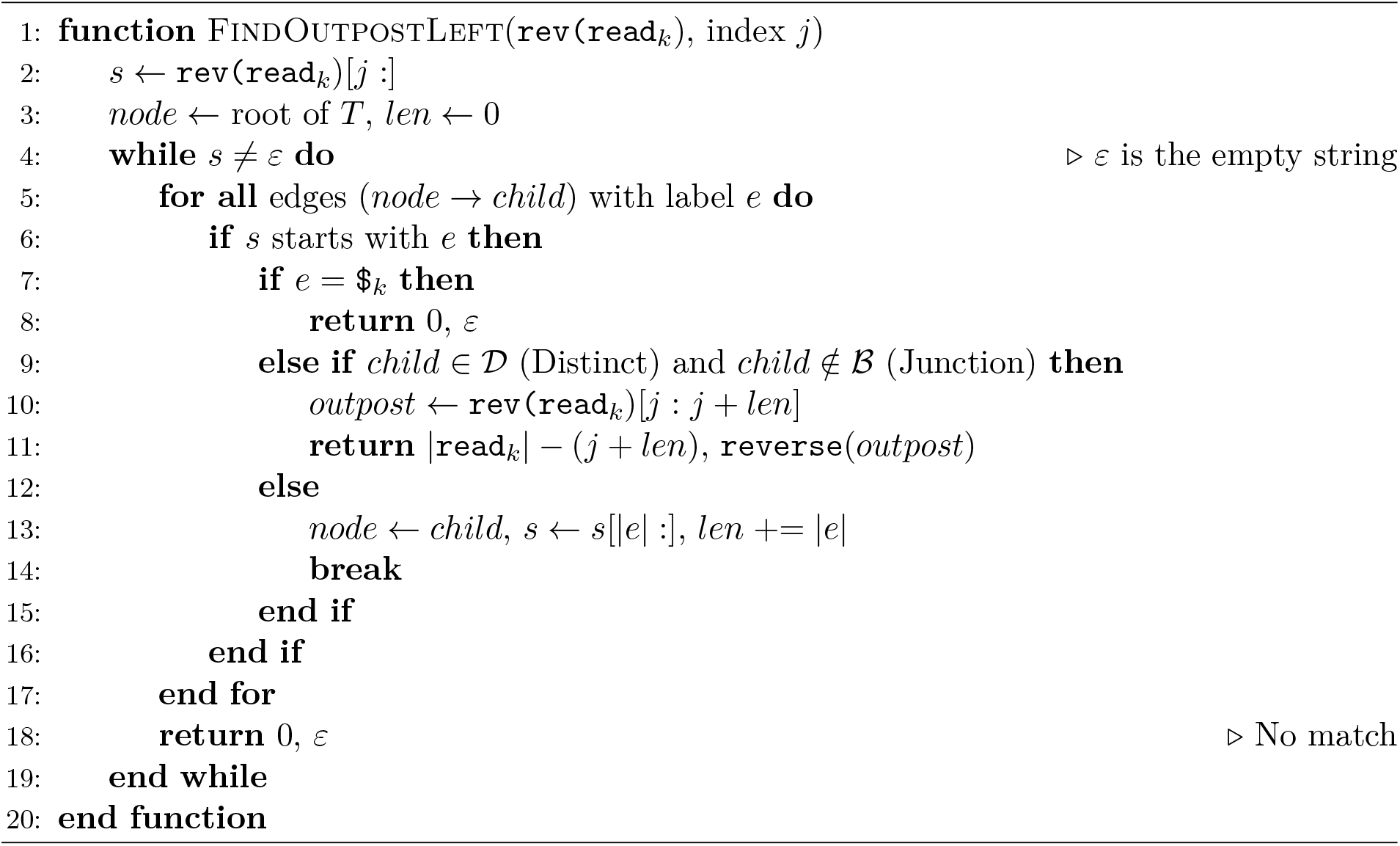

### 3.3 Identification of MUS

After building the generalized suffix tree 𝒯 (ℛ), we search it in a depth-first manner. This helps us find the MUSs within ℛ, using the outpost definitions and proposition 1 set up earlier. The collection of all identified MUSs is termed the `anchor set` and is denoted as *A*_ℛ_ (*w*). Our framework, inspired by [15], provides a way to assess the uniqueness of a word within ℛ.

Following proposition 1, a word *w* is consistent if its length from position *i* to *j* satisfies *j ≥* `Right`_*k*_ (*i*) and *i ≤* `Left`_*k*_(*j*), where `Right`_*k*_ (*i*) and `Left`_*k*_(*j*) are the outpost boundaries defined earlier. Building on this notion of consistency, we define the algorithmic `anchor set` *A*_ℛ_ (*w*). A word *w* = *D*[*i* : *j*], therefore, belongs to *A*_ℛ_ (*w*) if it meets the following three MUS-derived conditions:

1. **Consistency (***C*_1_): *w* must be consistent with respect to ℛ, ensuring it can be uniquely placed in the assembly.
2. **LMUS condition (***C*_2_): *w* cannot be shortened from the left without losing consistency. That is, while *D*[*i* : *j*] is consistent, its proper suffix *D*[*i* + 1 : *j*] is not: *j <* `Right`_*k*_ (*i* + 1) or *i* + 1 *>* `Left`_*k*_(*j*).
3. **RMUS condition (***C*_3_): *w* cannot be shortened from the right without losing consistency. That is, while *D*[*i* : *j*] is consistent, its proper prefix *D*[*i* : *j −* 1] is not: *i >* `Left`_*k*_(*j −* 1) or *j −* 1 *<* `Right`_*k*_ (*i*).

A word satisfying all three conditions is included in the anchor set *A*_ℛ_ (*w*). Algorithm 4 outlines the procedure for extracting *A*_ℛ_ (*w*) based on these conditions.

#### Algorithm 4

Compute *A*_ℛ_ (*w*) from `Right_k` and `Left_k`

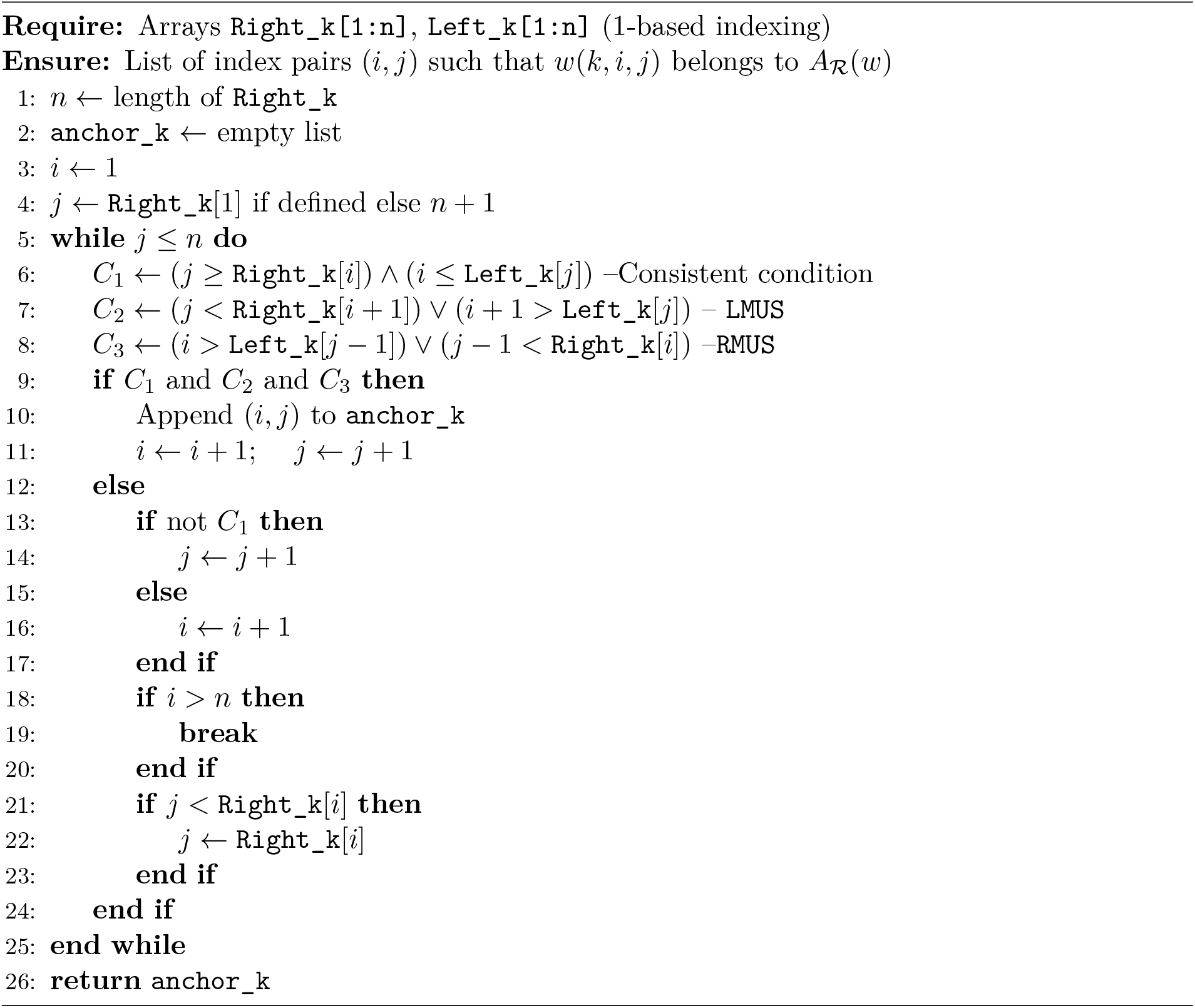

## 4. Empirical evaluation

### 4.1 Datasets and Experimental Setup

We tested the proposed framework on two genomes, *Escherichia coli K12* and *Humans*, which differ in size and repetitive DNA. Table 1 shows their main characteristics.

**Table 1:** Benchmark genomes used for MUS extraction.

| Genome | Num of Seqs | Total Size (Mb) | Repeats (%) |
| --- | --- | --- | --- |
| *E. coli K12* | 8,959 | 130.4 | 15 |
| Humans Chr11 | 5,626 | 84.0 | 45 |

All the analyses were done on a PowerEdge M620 (VRTX), equipped with 640.0 GB of RAM and powered by two Intel(R) Xeon(R) CPU E5-2670 v2 processors, operating at 2.50 GHz and running Ubuntu Server 22.04.5 LTS.

### 4.2 Performance Evaluation

#### 4.2.1 Runtime Assessment

We evaluated computational performance by analyzing execution time and peak memory consumption as a function of total sequence length. These metrics were delineated into two operational phases: suffix tree (ST) construction and Minimum Unique Substring (MUS) extraction. Given our use of a custom suffix tree implementation, empirical validation was paramount to confirm adherence to the theoretical linear time and space complexity (𝒪(*n*)). Tables 2, 3, and 4 show how the process scaled up across the two sets of data.

**Table 2:** Runtime and memory performance for the MUS extraction algorithm.

| Genome | Runtime (min) | Memory (GB) | MUSs |
| --- | --- | --- | --- |
| *E. coli k12* | 11.20 | 24.66 | 1,640,108 |
| Humans Chr11 | 8.38 | 13.59 | 970,845 |

**Table 3:**
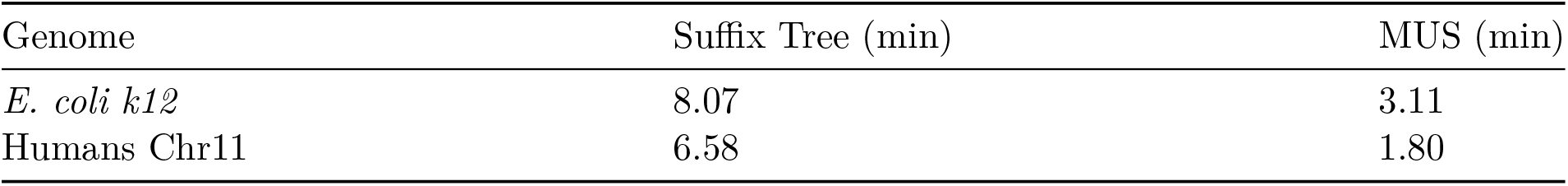
Runtime performance for each phase of the algorithm in minutes.

| Genome | Suffix Tree (min) | MUS (min) |
| --- | --- | --- |
| *E. coli k12* | 8.07 | 3.11 |
| Humans Chr11 | 6.58 | 1.80 |

**Table 4:**
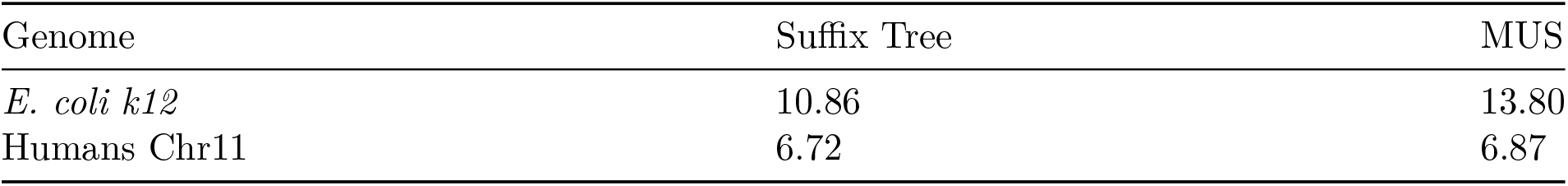
Memory performance for each phase of the algorithm in gigabyte (GB)

The benchmarks confirmed that both the construction and extraction phases scaled linearly with increasing input data. Processing the *E. coli* K-12 dataset (130.4 Mb) required approximately 11.2 minutes in total, comprising 8.07 minutes for ST construction and 3.11 minutes for MUS identification. In comparison, the human dataset was processed in approximately 8.40 minutes (6.58 minutes for ST; 1.80 minutes for MUS). Memory consumption exhibited a consistent linear trend: the *E. coli* dataset utilized a peak of 24.66 GB (10.86 GB for ST; 13.80 GB for MUS), while the human dataset required 13.59 GB (6.72 GB and 6.87 GB, respectively). These findings substantiate the 𝒪(*n*) efficiency of the pipeline, demonstrating that while total sequence volume is the primary driver of resource demand, secondary variations in performance are influenced by genomic architecture and local sequence redundancy rather than algorithmic scaling bottlenecks.

### 4.3 The Distribution of MUS Lengths

#### 4.3.1 MUS Length Variation Across Genomes

We looked at the length distribution of MUSs in each genome. Table 5 and Figure 5 show the MUSs distribution at different lengths.

**Table 5:** Summary statistics of MUS lengths across genomes.

| Genome | Mean Length (bp) | StD (bp) | Range (bp) |
| --- | --- | --- | --- |
| *E. coli K12* | 30.44 | 187.5 | 8-11337 |
| Human Chr11 | 36.08 | 222.98 | 7-9323 |

**Figure 5:**
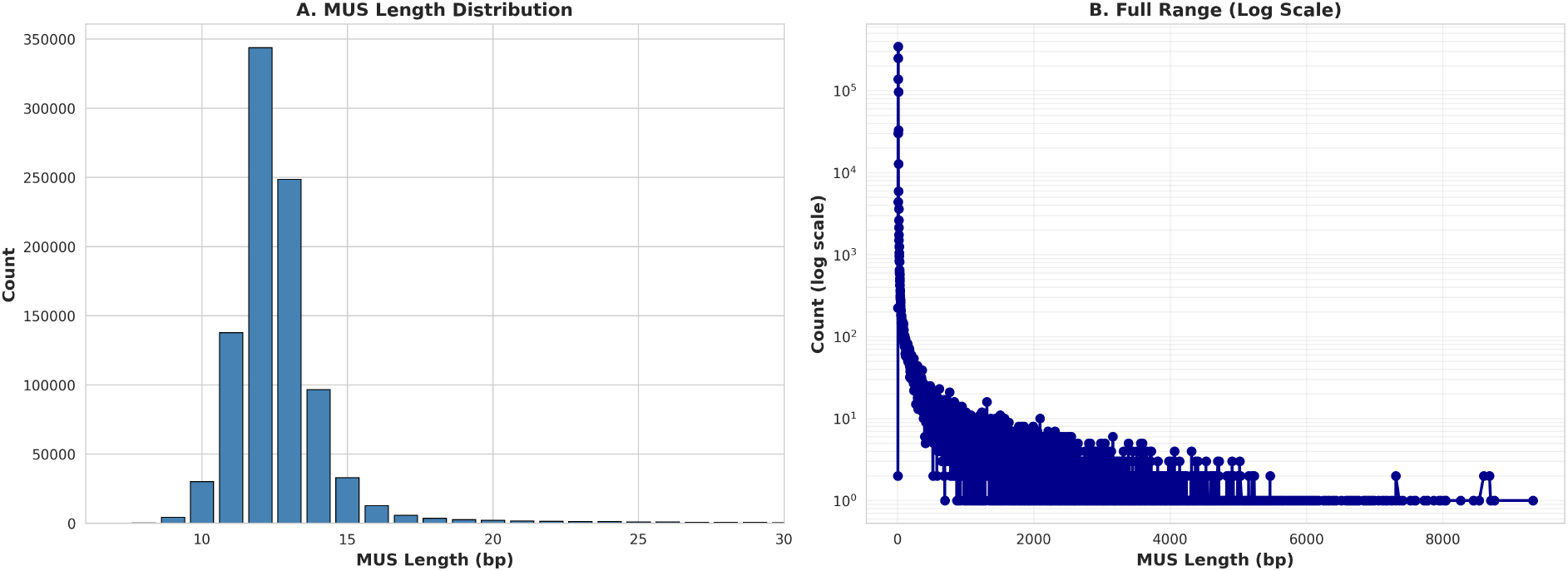
Distribution of MUS lengths in the human chromosome 11. (A) The Linear histogram shows higher peak at 12-13 bp with a broader spread (10-18 bp). (B) Log-scale view shows an extensive long tail of >8,000 bp with denser population around 100-5000 bp indicating the high complexity of mammalian genomes.

The length distributions of MUS reveal the differing genomic complexities of *E. coli K12* and Human Chr11. In *E. coli K12*, the distribution is tightly constrained, with 85% of MUSs falling between 11-13 bp (base pairs). It showed a 1417-fold dynamic range of 8-11337 bp (see Figure 5). This indicates a compact and high-complexity genome, where low repeat content (about 15%) allows the statistical floor for uniqueness to be reached almost immediately. In contrast, the Human Chr11 shows a wider range of MUS lengths (1332-fold dynamic range of 7-9323 bp), with a noticeable increase extending from 15 to 18 bp (see Figure 6). This trend toward longer MUSs is directly linked to the higher repetitive density in the human genome (around 45-50%). Because a MUS must be unique, any substring that begins within a repeat must extend until it crosses the boundaries of the repeat to obtain a unique flanking anchor. Therefore, while unique genomic regions produce shorter MUSs, repetitive regions yield longer MUSs that serve as context-aware markers of local complexity, adjusting their resolution without requiring manual parameter changes.

**Figure 6:**
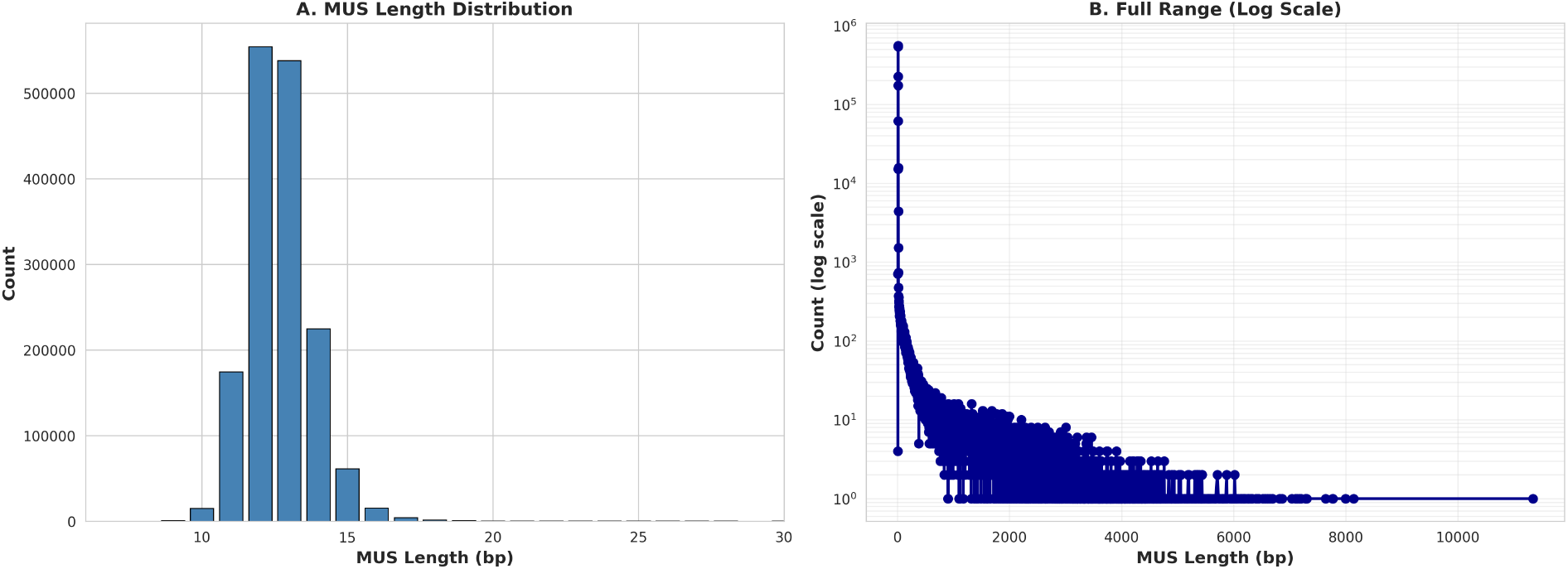
Distribution of MUS lengths in the *E*.*coli K12* genome. (A) The Linear histogram shows majority of MUS (>80%) concentrated at 10-15 bp. (B) Log-scale view shows the full dynamic range extending to 10,000+ bp. The variable-length distribution shows that no single fixed *k* can optimally represent the genomic complexity.

### 4.4 Comparison with Fixed *k*-mers

We evaluated the efficiency of information preservation and the inherent *k*-selection trade-off by benchmarking fixed-length *k*-mers against the MUS framework. Using the Human Chr11 dataset, *k*-mer spectra were generated (via Jellyfish) for *k* = 21, 31, 41, and 61 (see Table 6). As *k* was increased, the total number of distinct *k*-mers nearly doubled from 4.8 million to 9.9M (million), while the count of unique *k*-mers (singletons) rose sharply from 2.35M to 6.86M (see Figure 7). This expansion demonstrates that larger *k* values do not inherently reduce redundancy; instead, they produce ‘spurious uniqueness’ by shattering repetitive structures into unique subsequences simply because the *k*-mer length exceeds the local repeat unit.

**Table 6:** Summary Comparison of MUSs vs. Fixed *k*-mer Approaches

| Metric | MUSs | $k = 21$ | $k = 31$ | $k = 41$ | $k = 61$ |
| --- | --- | --- | --- | --- | --- |
| Total distinct tokens | 970,845 | 4,800,073 | 6,186,979 | 7,483,811 | 9,887,060 |
| Unique tokens (singletons) | 970,845 (100%) | 49.0% | 57.3% | 62.7% | 69.4% |
| Mean token length (bp) | 36.08 | 21 | 31 | 41 | 61 |
| Token redundancy (%) | 0.0 | 51.0 | 42.7 | 37.3 | 30.6 |
| Token reduction vs each $k$ (%) | — | 79.8% | 84.3% | 87.0% | 90.2% |
| Diploid coverage peak (×) | N/A | 36 | 35 | 35 | 34 |
| Error $k$-mers $\leq 5\times$ (%) | 0 (consistency-based) | 54.6 | 63.6 | 69.3 | 76.2 |
| Information content (bits/nt) | 0.5512 | 1.0079 | 0.7019 | 0.5405 | 0.3723 |
| MUS length mean (bp) | 36.08 | Fixed | Fixed | Fixed | Fixed |
| MUS length range (bp) | 7–9323 | — | — | — | — |

**Figure 7:**
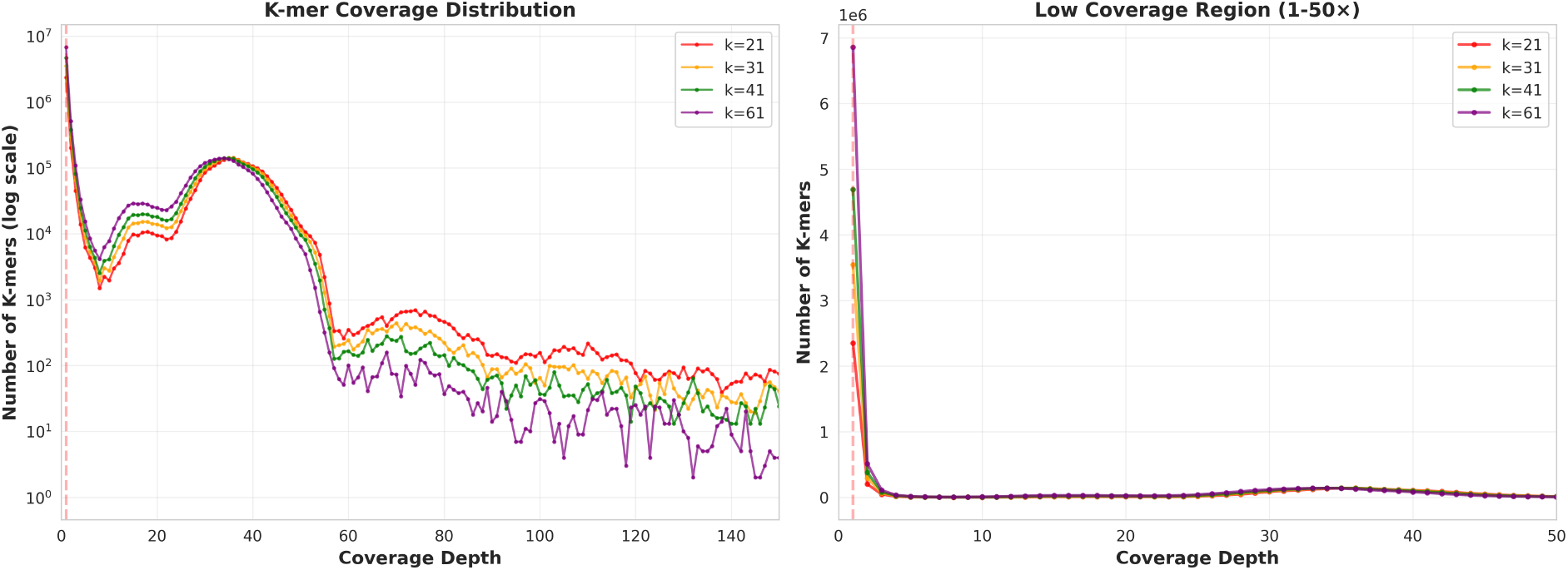
K-mer coverage distributions highlight the *k*-selection trade-off observed in Human Chromosome 11. The frequency histograms of *k*-mers at *k*= 21, 31, 41, and 61 show that while they converge at true genomic coverage ( 30-40x), there is a dramatic divergence at a coverage of 1. At this point, the counts of unique *k*-mers rises from 2.4M for *k*=21 to 6.9M for *k*=61. This illustrates that larger *k* values break down repeats into seemingly unique subsequences, rather than enhancing genomic coverage.

In contrast, the MUS approach achieved 100% unique position coverage with an average length of 36.08 bp, which is significantly more efficient than fixed-length approaches. For instance, even at *k* = 61, nearly twice the average MUS length, the *k*-mer approach attains only 69% unique coverage. By utilizing variable-length adaptation, MUSs maintain complete uniqueness while remaining compact. The inefficiency of the fixed-length approach is most evident at *k* = 61. Despite being 5.6 times longer than the average MUS, it achieved only 69% unique coverage, highlighting the superior sensitivity and resolution offered by the MUS framework.

## 5. Discussion

The results of this study show that minimum unique substrings (MUSs) are a theoretically sound and empirically validated, parameter-free alternative to fixed-length *k*-mers for representing genomic sequences. The results address the fundamental structural limitations of the fixed-*k* paradigm that are not amenable to parameter tuning. Also, these provide a direct pathway for assembly graph models in which node labels are derived from the data rather than fixed in advance.

Notably, the distribution of MUS lengths provides a direct high-resolution map of genomic complexity. In the compact *E. coli* K-12, with a repeat content of ∼15%, more than 80% of MUSs are located within a narrow range of a 10-13 bp window. This tightly constrained distribution means that uniqueness is achieved almost immediately at any locus. Also, the long tails (up to ∼11,000 bp in length) exist only at repeat boundaries. In such cases, MUSs must extend to cover the entire near-identical sequence to capture the unique flanking context. In contrast, the human chromosome 11, which has a much higher repeat density (∼45%), shows a broader primary distribution (mode ∼12 bp, with a marked shoulder extending to 18 bp) and a significantly longer tail extending beyond 8,000 bp. Importantly, both organisms show a similar mode, suggesting that the minimum threshold for uniqueness remains consistent across different genomic architectures. Nevertheless, the divergence observed in the upper tail is driven solely by the architecture of repetitive elements—specifically their density, length, and degree of similarity. Because these distributions are entirely inherent to the data, they require no parameter tuning to manifest. Instead, they provide a direct, quantitative signature of the genomic repeat landscape. Furthermore, the comparison with fixed-length *k*-mers reveals what we term the *k*-paradox.

This phenomenon represents a systematic failure of *k*-mer-based uniqueness to serve as a reliable proxy for genomic uniqueness. As *k* increases from 21 to 61 on human chromosome 11, the number of unique *k*-mers (singletons) nearly triples (from 2.35 million to 6.86 million) even though the same set of genomic positions is being represented. Notwithstanding, this expansion does not reflect improved genomic resolution. Rather, it is the fragmentation of repetitive sequences into substrings that are unique only in being longer than the local repeat unit. For example, a 61-mer across a boundary within a 40-bp tandem repeat is unique, but this uniqueness contains no additional biological information about the particular locus. Consequently, this uniqueness is a statistical artifact of the window size. We refer to this phenomenon as spurious uniqueness, and show that it accounts for an increasing fraction of the singleton pool as *k* increases.

In contrast, the MUS framework entirely escapes this paradox. Because an MUS extends only until genuine uniqueness is achieved, every marker in the anchor set corresponds to a single, biologically meaningful genomic locus. Consequently, the framework achieves 100% positional coverage with 7.1*×* fewer tokens than the *k*=61 representation while remaining zero-redundant by construction. The read-consistent extension of MUS theory formalized in this work provides the theoretical foundation for applying these properties to de novo genomic sequence analysis. Prior MUS theory was restricted to single, known sequences, rendering it inapplicable to fragmented sequencing datasets. The *Superstring*(*w*) consistency criterion bridges this gap. By requiring that all reads containing a putative MUS agree on its placement within a unique minimal superstring, this framework seamlessly accommodates fragmented data. Crucially, this is not a heuristic relaxation of uniqueness but a mathematically rigorous extension that fully preserves the minimality guarantees of the single-string definition. The outpost node framework translates this theory into an efficient computational solution. Outposts act as topological anchors within the Generalized Suffix Tree, marking the exact nodes where a subtree transitions from a branching (repetitive) to a non-branching (unique) structure. This topological definition enables the algorithm to precisely localize MUS boundaries in 𝒪(*n*) time without a reference sequence.

Both the suffix tree construction and the MUS extraction scale linearly, confirming the tractability of the framework at the whole-genome scale. We observe a difference in performance between the two stages: The *E. coli* dataset has a larger share of the total runtime for MUS extraction due to the full outpost traversal. This is the expected behavior. Tree building involves one-pass insertion, whereas uniqueness testing examines all suffix paths. This trade-off is easily justified. We provide an 𝒪(*n*) guarantee of exact MUS identification, a necessary prerequisite for the correctness of all downstream analyses, at a theoretically optimal asymptotic cost.

## 6. Conclusion

This study presents Minimum Unique Substrings (MUS) as a theoretically rigorous and biologically meaningful alternative to fixed-length *k*-mers for genomic sequence representation. The three contributions of this work, including the read-consistent formalization of MUS uniqueness, the outpost-based 𝒪 (*n*) extraction algorithm, and the empirical characterization of MUS distributions across genomic architectures, collectively establish MUSs as a principled foundation for parameter-free sequence analysis.

The key finding is that MUSs achieve 100% positional uniqueness at substantially lower token counts and shorter mean lengths than any tested fixed *k*-mer configuration. This is not an incremental improvement over *k*-mers but a consequence of a qualitatively different approach to sequence representation. Where *k*-mers impose a uniform resolution on a heterogeneous genome, MUSs allow resolution to emerge from the data, adapting to local sequence complexity through the minimality constraint. The *k*-paradox demonstrates that fixed-length uniqueness is not equivalent to genomic uniqueness. It reveals that the inflation of apparent unique tokens with increasing *k* is a representational artifact rather than a biological signal.

MUSs, on the other hand, are immune to this artifact by construction. The length distributions observed across *E. coli* K-12 and human chromosome 11 validate the theoretical prediction that MUS lengths encode genomic architectural complexity. The shared modal length (∼12 bp) across both genomes establishes a universal minimum threshold for genomic uniqueness in these organisms. Also, the divergence in the upper tails directly quantifies the additional information required to span the repeat landscapes of each genome. This property has immediate utility in genomics sequence analysis. Thus, MUS as graph nodes in assembly graphs, and their length distributions, can serve as parameter-free metrics of repetitive content, indicators of structural variation at specific loci, and genomic complexity fingerprints for alignment-free comparative analysis. For genome assembly specifically, the duality between MUSs and maximum repeats – which guarantees that every MUS node in an assembly graph lies at the boundary of a repeat – means that an assembly graph constructed from MUS nodes cannot collapse distinct repeat copies. Repeat resolution is a structural property of such a graph, not a post-hoc heuristic.

The formalization of read-consistent MUSs and the outpost algorithm that extracts them provides the first principled, parameter-free definition of assembly graph node labels that directly encodes the uniqueness structure of the underlying genome. The framework is immediately applicable in three domains. In repeat characterization, MUS length distributions offer a biologically interpretable, parameter-free signature of genomic complexity that complements and extends existing repeat-annotation pipelines. In alignment-free comparative genomics, read-consistent MUSs provide position-specific anchors for cross-genome comparison that do not depend on alignment or any external reference. In the assembly graph, the anchor set produced by this framework provides the node labels for a graph model in which parameter sensitivity is eliminated by design. Together, these applications represent a substantive shift from fixed-length to adaptive sequence representation – a shift that is not merely pragmatic but is grounded in the formal structure of genomic sequence uniqueness.

## 7. Limitations and Recommendations

The empirical validation of the MUS framework across both prokaryotic and eukaryotic datasets confirms the soundness of the underlying theoretical model. The limitations identified here pertain to the current implementation rather than to the model itself. Crucially, each limitation is technically bounded and admits a well-defined resolution.

The primary implementation constraint is the memory overhead of the uncompressed Generalized Suffix Tree. Both tree construction and MUS extraction operate in 𝒪 (n) time. However, the space requirement of 𝒪 (*n*) words–rather than 𝒪 (*n*) bits–imposes a high constant-factor cost. This overhead becomes prohibitive for genomes that substantially exceed the tested range of approximately 130 Mb. We deliberately adopted the uncompressed tree to ensure the mathematical exactness of MUS boundary identification. This exactness is a prerequisite for the correctness of the read-consistent uniqueness checks. Fortunately, the path to resolution is well-characterized. Replacing the uncompressed tree with a succinct data structure preserves the 𝒪*n*)-time extraction guarantee. Suitable alternatives include a compressed suffix array, an FM-index, or a wavelet tree. This modification reduces the required space to 𝒪 (*n*) bits. As a result, the optimized framework is projected to handle complete human and polyploid genomes within a standard 128-256 GB RAM envelope. This transition does not affect any theoretical property or algorithmic correctness result presented in this paper.

Future work will proceed along four distinct directions. First, we will integrate succinct indexing structures to achieve true whole-genome scalability. Second, we will construct and evaluate a complete *v*MUS-dBG assembly pipeline. This pipeline will incorporate MUS nodes as graph node labels and will be benchmarked against existing assembly tools. Third, we will extend the read-consistent MUS framework to polyploid and metagenomic settings. Finally, we will apply MUS-based anchor sets to read mapping and structural variant detection. In these domains, the parameter-free nature of MUS boundaries is expected to offer significant advantages over traditional, fixed-length seed-selection methods.

### Data and Source Code

git@github.com:fandrew19/vMUS-dBG-Assembler.git

